# Mud, microbes, and macrofauna: seasonal dynamics of the iron biogeochemical cycle in an intertidal mudflat

**DOI:** 10.1101/2020.05.18.100800

**Authors:** Jacob P. Beam, Sarabeth George, Nicholas R. Record, Peter D. Countway, David T. Johnston, Peter R. Girguis, David Emerson

## Abstract

Microorganisms and burrowing animals exert a pronounced impact on the cycling of redox sensitive metals in coastal sediments. Sedimentary metal cycling is likely controlled by seasonal processes including changes in temperature, animal feeding behavior due to food availability, and availability of organic matter in sediments. We hypothesized that the iron biogeochemical cycle and associated sedimentary microbial community will respond to seasonal changes in a bioturbated intertidal mudflat. In this study, we monitored the spatiotemporal dynamics of porewater and highly reactive solid phase iron with the corresponding prokaryotic and eukaryotic sedimentary microbial communities over one annual cycle from November 2015 to November 2016. Continuous and seasonally variable pools of both porewater Fe(II) and highly reactive iron (Fe_HR_) were observed throughout the season with significant increases of Fe(II) and Fe_HR_ in response to increased sediment temperature in summer months. Maximum concentrations of Fe(II) and Fe_HR_ were predominantly confined to the upper 5 cm of sediment throughout the season. Iron-oxidizing and -reducing microorganisms were present and stable throughout the season, and exhibited strong depth-dependent stratification likely due to availability of Fe(II) and Fe_HR_ pools, respectively. Otherwise, the community was dominated by Deltaproteobacteria, which are involved in sulfur and potentially iron cycling, as well as Gammaproteobacteria and Bacteroidetes. The microbial community was relatively stable throughout the seasonal cycle, but showed strong separation with depth, probably driven by changes in oxygen availability and organic matter. The relative abundance of diatoms revealed a noticeable seasonal signature, which we attribute to spring and fall blooms recorded in the sediments. Macro-, meio, and microfauna were detected throughout the season with some seasonal variations that may influence sedimentary iron transformations by active microbial grazing. The seasonal dynamics of the sedimentary iron cycle are controlled by numerous, interdependent processes, with macrobiota-microbiota relationships and depth stratification comprising primary components. Deciphering these processes in natural ecosystems is essential to understand how they might respond to future environmental perturbations, such as anthropogenic nutrient release to coastal systems.

## Introduction

Coastal marine sediments are very active and dynamic environments that connect the flux of nutrients and energy from weathered land masses to marine ecosystems. Continentally-derived energy and nutrient sources are utilized and recycled by pelagic and benthic organisms, which act as key biogeochemical agents in coastal ecosystems. Biogeochemical cycles in coastal sedimentary ecosystems can fluctuate on hourly to seasonal time scales, and represent dynamic gradients in space and time influenced by a myriad of biogeophyiscal processes such as riverine discharge, tidal cycles, seasonal stratification, animal behavior, and microbial activity (Kristensen and Kostka, 2005; Bouma et al., 2009). In coastal sediments, sediment-dwelling animals have played a primary top-down control on biogeochemical processes over the last ~500 million years (Thayer, 1979; Meysman et al., 2006; Tarhan et al., 2015). These ecosystem-engineering animals actively burrow in sediments and irrigate their burrows with the overlaying bottom water, supplying oxygen and nutrients deep into oxygen-free (i.e., reduced) sediments. Their burrow walls are hotspots for microbial redox cycling where steep geochemical gradients drive the rapid cycling between reduced and oxidized chemical species (Reichardt, 1986; Aller, 1994; Fenchel, 1996; Brune et al., 2000; Kristensen and Kostka, 2005; Bertics and Ziebis, 2009). Their complex burrowing and irrigating behavior essentially increases the exchange of solutes between benthic and pelagic systems (Aller, 1982), and are thus of primary importance to benthic-pelagic coupling and ecosystem function (Griffiths et al., 2017).

Iron is an essential electron donor and acceptor in microbial metabolism (Emerson et al., 2010; Melton et al., 2014) and a universally important micronutrient source. Numerous, interdependent biological, chemical, and physical processes control the redox cycling of iron in sedimentary environments. Iron is transported to coastal ecosystems by rivers where it settles in sediments and is intensely [re]cycled (Canfield, 1989; Canfield et al., 1993). Sediment-dwelling animals act in concert with diverse microbiota in coastal sediments to regulate iron biogeochemical transformations (Thibault de Chanvalon et al., 2016; Beam et al., 2018; van de Velde et al., 2020). These macrobiota-microbiota relationships result in labile pools of dissolved and highly reactive iron that contribute to important ecosystem processes such as formation of bioavailable iron and subsequent transport to the water column (Elrod et al., 2004; Severmann et al., 2010) and inhibition of porewater hydrogen sulfide accumulation (Seitaj et al., 2015). These labile pools of iron provide a continuous replenishment of reduced and oxidized iron for iron-oxidizing and iron-reducing microorganisms, respectively (Canfield, 1989; Canfield et al., 1993; Beam et al., 2018). In addition to these direct biological factors, the sedimentary iron biogeochemical cycle may also respond to fluctuations in temperature and organic matter transport from primary productivity in the water column. Based on this knowledge, we have a reasonable understanding of some of the major drivers of coastal iron biogeochemistry. However, an important aspect that is missing from the current view of the sedimentary iron biogeochemical cycle is how seasonal changes, particularly with regard to the composition and function of sedimentary microbial communities, impact iron biogeochemistry. In many ecological systems, sub-annual seasonal dynamics are as important or can even outweigh year-to-year variation (Staudinger et al., 2019). Thus, it is essential to understand the seasonal dynamics of these interconnected processes, in order to predict how they might respond to effects of anthropogenic climate change in coastal sedimentary ecosystems such as increased temperature, declines in oxygen, and eutrophication (Doney, 2010).

In this study, we tracked the spatial and temporal dynamics of the iron biogeochemical cycle over one seasonal cycle (2015-2016) in a bioturbated intertidal mudflat to determine changes in the reduced porewater Fe(II) and highly reactive Fe (Fe_HR_) mineral pools. The shifts in the sedimentary prokaryotic and eukaryotic microbial communities were also spatiotemporally tracked with 16S and 18S rRNA gene sequencing, respectively. These data suggested a relatively continuous and dynamic pool of both Fe(II) and Fe_HR_ over the season, and statistically significant increases of both pools with increasing temperature. The microbial communities were relatively stable over the annual cycle with some pronounced seasonal and spatial variability, possibly due to interactions with water column processes such as phytoplankton blooms. Seasonal and spatial shifts in the sedimentary pools of porewater and solid phase iron are controlled by the intricate interaction between sediment-dwelling animals, sedimentary microbial communities, and water column processes that are a result of complex interconnected processes.

## Materials and Methods

### Sampling site

An intertidal mudflat (the Eddy, 43.994827, −69.648632) located within the Sheepscot River Estuary Maine, United States of America was the subject of the current study (Figure 1) (Stickney, 1959). As defined by Stickney (1959) the Eddy would be in the middle of the upper and lower portions of the Sheepscot Estuary, and the large observed seasonal variations in salinity and temperature would classify this mudflat as upper estuary receiving significant influence from brackish and/or freshwater riverine input (Stickney, 1959). From November 2015-November 2016 sediment cores were sampled (n=11; no data for August and October 2016) for porewater and solid phase geochemistry in combination with microbial community structure (n=10) at 1 cm depth intervals up to a depth of 10 cm. We took care not to take monthly samples in areas that had been previously disturbed by sampling the previous month, and by local worm and shellfish harvesters as evinced by visible digging tracks in the surface sediments (Fig. 1A). A wooden post and a washed-up bundle of lobster traps served as a reference point for sampling throughout the season (Fig. 1B). At high tide, this area of the mudflat was covered with approximately one meter of water, dependent on tidal amplitude. At low tide, we successively took samples in the same horizontal distance from shore to reduce possible longitudinal effects on the mudflat macrofaunal populations. This mudflat lies adjacent to a tidally flushed salt marsh to the west, and faces south-southwest to the main stem of the Sheepscot River. The salinity and temperature varied throughout the season, but the overlying water was never frozen in the winter months of 2015-2016. All physicochemical data are located in Table S1.

**Figure 1.**
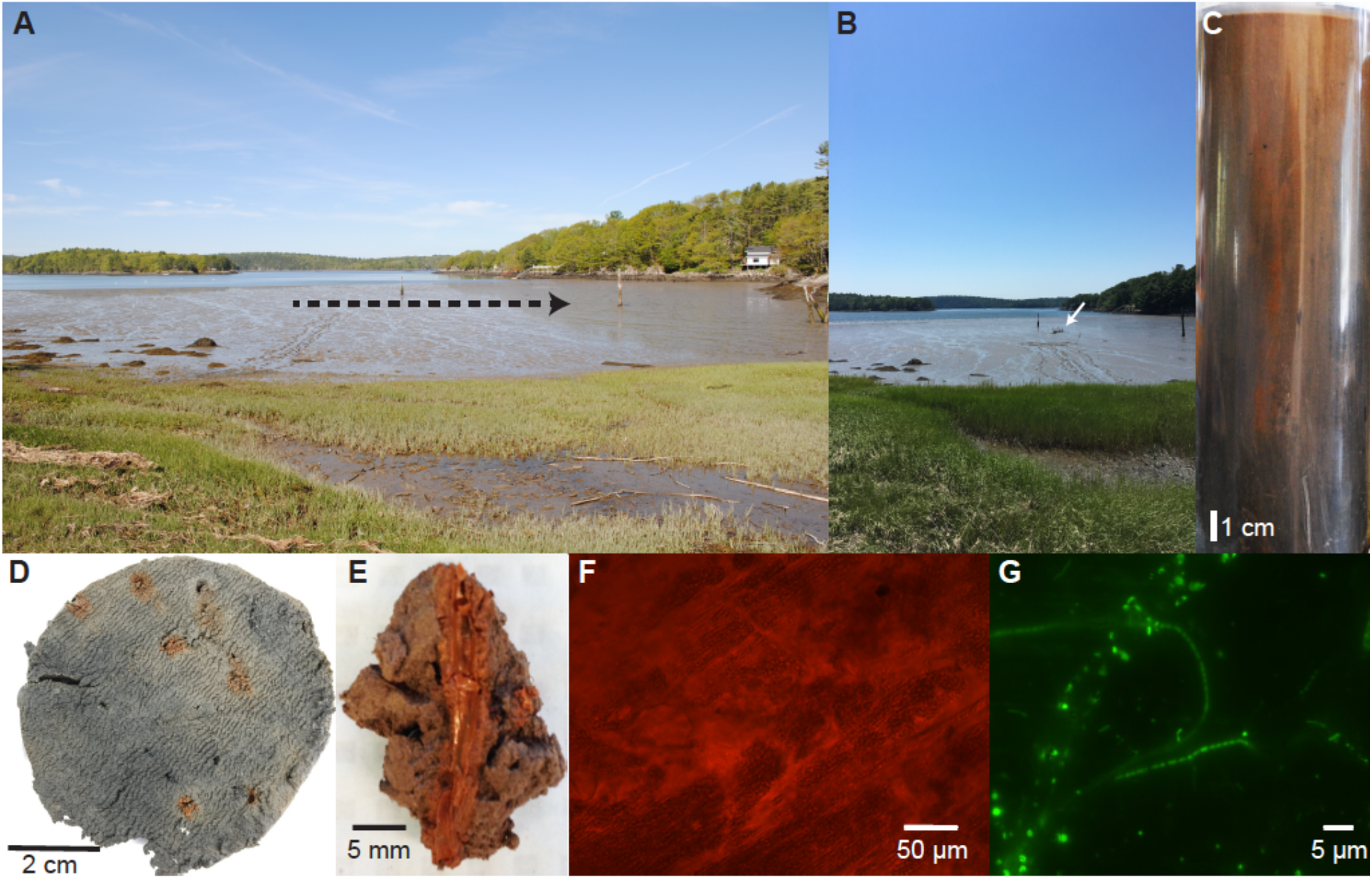
A photomontage of images over different time and spatial scales from seasonal sampling at “The Eddy” (Maine, USA; 43.994827, −69.648632). A landscape view of the mudflat at low tide (**A**) (26 May 2016) with dashed arrow line depicting the progression of sampling throughout the annual cycle (tracks created from sampling are visible at the position distal to the arrowhead). A representative core (**C**) (18 November 2015) from “The Eddy” of ~20 cm of total sediment showing deep mixing by macrofauna up to ~15-17 cm in depth by visible iron oxides, then the sediments become bluish-grey to black. A one-centimeter core slice from 5 cm depth (**D**) showing characteristic iron oxide lined worm burrows most likely created by the common polychaete *Nereis diversicolor* and/or the intertidal inhabiting hemichordate *Saccoglossus kowalevskii*. Close up image of a dissected iron oxide lined burrow (**E**) (28 December 2015) and light microscope image of burrow lining at 400 x magnification (**F**). Fluorescence microscope image of DNA stained microorganisms inhabiting the iron oxide burrow lining (**G**) (8 October 2015). Filamentous *Beggiatoa* spp. (thick filament) And putative ‘cable bacteria’ (thin filaments) were commonly observed throughout the season.

### Porewater Fe(II) geochemistry

Sediment cores (approximately 15-20 cm in depth) from the Eddy were taken with a Plexiglas core barrel (inner diameter, 7.5 cm) that had predrilled 4 mm holes drilled (sealed with electrical tape before sampling) at 1 cm intervals for Rhizon CSS samplers with a 5 cm length and 0.15 μm pore size (Rhizosphere Research Products, Wageningen, Netherlands) (Seeberg-Elverfeldt et al., 2005). Porewaters were extracted by inserting Rhizons into the predrilled holes starting from the top of the core and working down to the final depth interval (9-10 cm). Sterile 10 mL syringes were used to pull negative pressure on the Rhizons, and the plunger was held in place for 1-2 hours with a wooden block. After ~10 mL of porewater had been extracted, the Rhizons were removed, and the porewater was dispensed into a sterile 15 mL tube in a glove bag containing an atmosphere of N_2_/H_2_ (95/5 %) or under ambient atmospheric conditions. Ferrous iron [Fe(II)] was immediately preserved by pipetting 250 μL of pore water into 250 μL of Ferrozine buffer (10 mM Ferrozine in 50 mM HEPES buffer, pH=7) (Stookey, 1970; McBeth et al., 2011). We determined that rapid preservation of porewater Fe(II) with Ferrozine under ambient atmospheric conditions was no different than preserving it in a glove bag, thus half of the samples were sampled under ambient atmospheric conditions. The Ferrozine-porewater solution was measured at 562 nm on a Multiskan MCC plate reader (Thermo Fisher) and absorbance values were calculated to concentrations by plotting them on a standard curve of known Fe(II) concentration solutions ranging from 0-1,000 μM prepared from a 10 mM solution of FeCl_2_ in 0.1 N HCl.

### Solid phase highly reactive Fe (Fe_HR_) geochemistry

Sediment cores used for porewater extraction were extruded and sliced into 1 cm sections with a brass wire and dried in an oven at 70 °C for 24 hours. Highly reactive iron oxides (ferrihydrite and lepidocrocite as well as iron monosulfides that were oxidized during this process) were extracted for 48 hours in a 50 mL falcon tube with 10 mL of hydroxylamine-HCl (1 M) in 25 % acetic acid (w/v) (Poulton and Canfield, 2005). The total extracted Fe was measured with a Ferrozine solution as described above, except the samples were diluted 1:100 in distilled water due to high absorbance values of undiluted samples.

### Sediment DNA extraction

Sub-cores were sampled simultaneously with porewater extraction using 1 mL syringes with cut off tips. The 1 mL syringe corers were inserted into pre-drilled 8 mm holes at a right angle to the Rhizon sampling holes, and gently pressed into the side of the core while drawing back on the plunger to remove approximately 0.2-0.5 cc of sediment. The 1 mL sub-cores were then frozen at −80 °C until DNA extraction. DNA was extracted from ~0.1-0.2 g of wet sediment using the DNeasy PowerSoil Kit (Qiagen) with a modification of the removal of 200 μL of the bead solution and addition of 200 μL of 24:25:1 phenol:chloroform:isoamyl alcohol (Sigma Aldrich); the manufacturers protocol was then followed for the rest of the extraction, with the exception of the DNA column elution step, which was eluted with 100 μL of molecular grade water instead of the elution buffer from the kit. The DNA was quantified using a Qubit 3.0 fluorometer with the Qubit dsDNA quantification kit and protocol (Thermo Fisher Scientific).

### 16S and 18 rRNA gene sequencing and analysis

The 16S rRNA gene was amplified from 1 cm interval DNA extractions with the 515F/926R primer pair and the 18S rRNA gene was amplified with the E572F/E1009R primer pair at the Integrated Microbiome Resource (Dalhousie University, Halifax, Nova Scotia, Canada). Paired end reads were assembled and classified using a mothur (Schloss et al., 2009) pipeline from the mothur MiSeq standard operating procedure (Kozich et al., 2013; Scott et al., 2017). Operational taxonomic units were clustered at either 85 % or 97%, which generated 12,437 and 351,057 OTUs, respectively. The 85% identity cutoff, which corresponds roughly to the class-order taxonomic level (Yarza et al., 2014), was chosen for comparative analysis to limit the number of clusters generated for comparisons and lend clarity to data representations. Relative abundances were utilized to build a heatmap in RStudio (RStudio Team) of the top OTUs that were greater than 0.5% abundance in one sample.

The 18S rRNA gene data was processed using mothur (v1.43) following the mothur MiSeq SOP data analysis pipeline (Kozich et al. 2013), but referencing the pR2 database (v4.12) to assign eukaryote taxonomies (Guillou et al., 2013). Eukaryote OTUs were determined at a level of 97% sequence similarity (Hu et al., 2015; Pasulka et al., 2019), which is less conservative than some thresholds proposed for rRNA genes (Caron et al., 2009), and more conservative than others (Callahan et al., 2017; Edgar, 2018). Counts of eukaryotic lineages of interest (i.e., diatoms, polychaetes, molluscs, nematodes, and protozoa) were filtered from the taxonomic data and used to calculate relative abundances for all samples across the season.

### Ecostates

One way to organize and find signals in complex ecological communities is to cluster them into coherent “ecostates” and then to map these ecostates back onto time and space to highlight patterns (Record et al., 2017). Each ecostate represents a consortium of taxa that clusters together consistently throughout the sampling program, as well as the respective relative abundances of taxa (O’Brien et al. 2016). For this analysis, samples were clustered using average linkage hierarchical clustering of the Bray-Curtis dissimilarity as implemented in the R package vegan (Oksanen, 2019). The clustering algorithm used bacterial and archaeal operational taxonomic units (OTUs) clustered at 85 % identity (n=12,437 OTUs), since, as discussed above, this identity was a compromise between capturing overall community diversity without overwhelming the analysis with a large number of OTUs. Ecostates were then numbered and mapped back onto the spatiotemporal sampling grid in R Studio (R Studio Team).

### Data availability

The pore water and solid phase sedimentary geochemical data can be found in the Biological and Chemical Oceanography Data Management Office (BCO-DMO) database under the DOI: 10.1575/1912/bco-dmo.737962.1. The 16S and 18S rRNA gene raw read data can be found under the Bioproject ID PRJEB36098. All sedimentary geochemical and DNA data can also be found in Table S1; all geochemical and DNA profile plots in the main text can be reproduced from this data set.

## Results and discussion

### Seasonal porewater dissolved ferrous iron [Fe(II)] pools

The dissolved ferrous iron [Fe(II)] in porewaters exhibited both spatial and temporal variability over the sampling period, which was approximately every month during the 2015-2016 annual cycle (Fig. 2A, B). Porewater Fe(II) concentrations peaked at a maximum average concentration of approximately 120 μmol L^−1^ at a depth interval between 3-4 cm, which is a consistent observation with many porewater Fe(II) profiles from coastal and continental shelf systems (Aller, 1980; Canfield, 1989; Hines et al., 1991; Thamdrup et al., 1994; Severmann et al., 2010), which are more often than not, bioturbated. This peak in porewater Fe(II) is consistent with the activity of iron-reducing microorganisms at this depth interval throughout the season. Throughout the season, the porewater Fe(II) concentration maxima remained in the top 5 cm of sediment, and migrated only once to a depth of 6.5 cm in September (Figure 2B). These Fe(II) porewater maxima suggest that the most active iron cycling zone occurs within the upper 5 cm of sediment. Concentrations of Fe(II) decreased to approximately a seasonal average of 100 μmol L^−1^ at the maximum depth interval sampled between 9-10 cm (Fig. 2A). The decline in Fe(II) with depth is likely due to the reaction with hydrogen sulfide produced by sulfate-reducing bacteria in sediments (Jørgensen, 1982) as well as around macrofaunal burrow walls (Bertics and Ziebis, 2010), which results in the precipitation of iron sulfides in sediments (Berner, 1984). There were consistent pools of porewater Fe(II) over the annual cycle, which exhibited large spatiotemporal changes (Fig. 2B). The large variability over space and time could be due, in part, to the heterogeneity of natural populations of macrofauna and microorganisms in the mudflat, which we attempted to minimize by sampling cores within a consistent region over the course of the study (Figure 1). Other studies have shown consistent variability in porewater Fe(II) with space and time (Hines et al., 1991; Thamdrup et al., 1994). The abundance and activity of macrofaunal populations likely exerts a primary control on porewater Fe(II) concentrations, due to their sediment mixing, irrigating, and feeding behavior (Aller, 1994).

**Figure 2.**
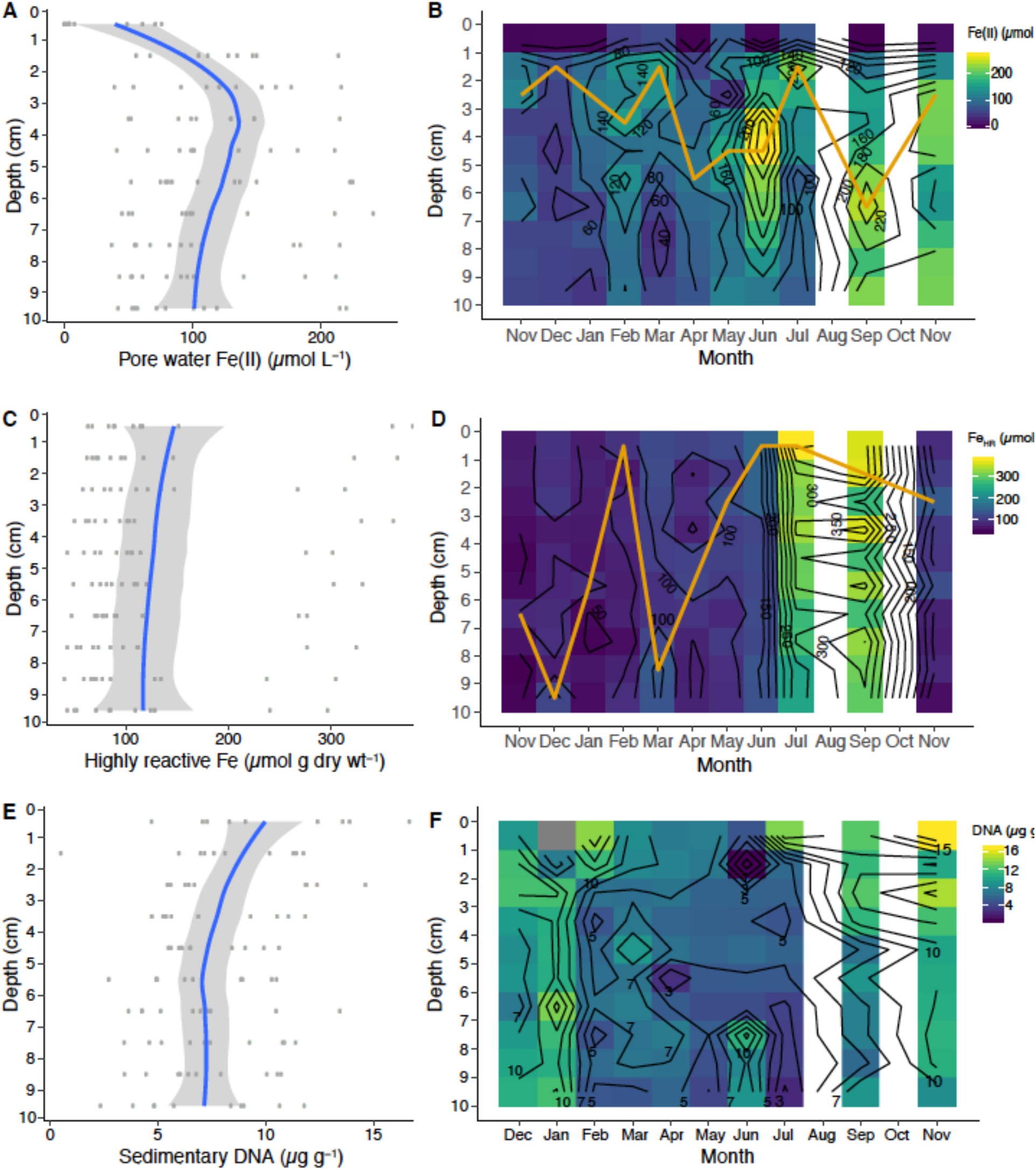
Porewater Fe(II), highly reactive Fe concentrations, and sedimentary DNA profiles from averages across depths and season (**A**, **C, E**) and over the annual seasonal cycle from Nov. 2015 to Nov. 2016 (**B**, **D, F**). Blue lines represent mean concentrations calculated from the individual seasonal data points (dark grey circles) and light grey shaded area is the 95 % confidence interval. The orange lines in **B** and **D** represent maximum iron concentrations.

Generally, the porewater Fe(II) concentrations increased with rising temperature (Fig. 3A). This might be indicative of elevated sedimentary carbon mineralization in summer months by iron-reducing bacteria (Canfield, 1989; Thamdrup et al., 1994). The increases in porewater Fe(II) with elevated temperatures between the months of February and July has also been observed in bioturbated sediments in the Great Bay New Hampshire, USA (Hines et al., 1984). The supply of terrigenous organic matter to this mudflat occurs mainly in the spring that corresponds with the thawing land mass and riverine run-off (Stickney, 1959). Organic matter may also originate by the density settling and/or active transport of large and small phytoplankton into the sediment by irrigating macrofauna (Christensen et al., 2000). The high mobility and turnover of porewater Fe(II) due to the behavior of irrigating animals influencing microbial iron cycling processes. This relationship likely explains the large variance of the porewater Fe(II) pools over the season, which is overall influenced by seasonal variations in sediment temperature (Figure 3A) and perhaps availability of macrofaunal food sources such as phytoplankton (discussed in detail below).

**Figure 3.**
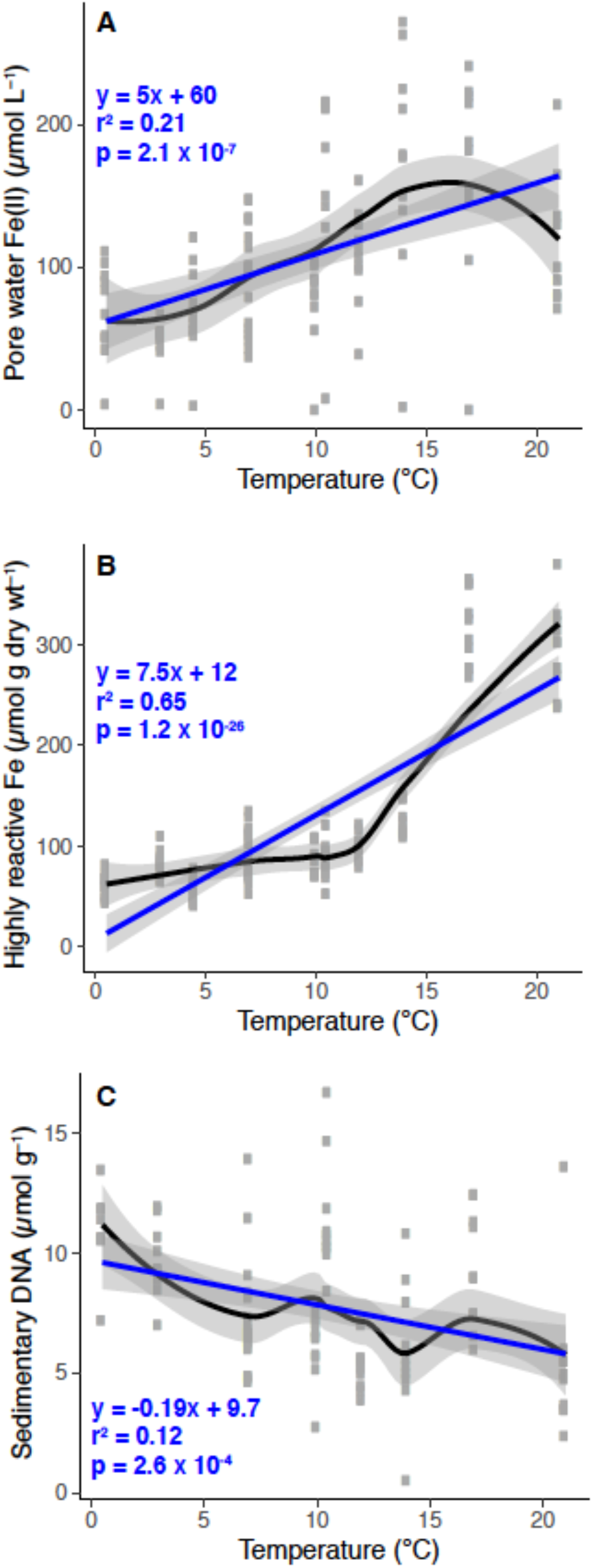
Porewater Fe(II) (**A**), highly-reactive Fe (**B**), and sedimentary DNA (**C**) pools as a function of temperature over the annual cycle at “The Eddy”. Individual data points are grey for each sampling event over the season. Black and blue lines are a loess smooth and a generalized linear model fit, respectively and grey shading represents the 95 % confidence interval for each line fit. The linear model statistics and equation are in blue text.

### Seasonal highly reactive Fe (Fe_HR_) pools

Similar to measurements of Fe(II), there was a continuous and variable pool of highly reactive Fe (Fe_HR_) with respect to depth over the seasonal cycle (Fig. 2C, D). The concentration of Fe_H_R declined slightly with depth (Fig. 2C) perhaps due to the interaction with hydrogen sulfide, which was not detected in these bioturbated sediments. The most significant change in the Fe_HR_ pool was in the warmer months (July and September) into the summer and early fall, with a nearly two-fold increase in the Fe_HR_ pools compared to the other months over the season (Fig. 2D). We hypothesize that this change is due to several key biogeochemical processes, which are controlled by the master variable, temperature, over the seasonal cycle. Changes in irrigation rates that likely occur throughout the season (Fang et al., 2019) will have profound effects on the exchange of solutes as well as the control of the pools of sedimentary Fe_HR_—when irrigational activity increases, concomitant microbial oxygen consuming process, such as microaerobic bacterial iron oxidation and aerobic carbon mineralization increase. When irrigation activity is low, the pools of sedimentary Fe_HR_ are depleted in the winter months. This is likely due to interaction Fe(II) with hydrogen sulfide produced by sulfate reducing microorganisms, resulting in the formation of more iron sulfides. An increase in the pool of Fe_HR_ in the summer has several important contributions to ecosystem function: 1) it prevents build-up of pore water H_2_S similar to a proposed “firewall” mechanism in seasonally hypoxic coastal areas (Seitaj et al., 2015), 2) it potentially enhances iron flux from the sediment by stimulating iron reduction (Severmann et al., 2010), and 3) provides fresh and reactive substrates for microbial iron reduction, which also effectively inhibits and/or decreases sulfate reduction in bioturbated sediments (Froelich et al., 1979; Canfield, 1989).

### Seasonal sedimentary DNA pools

The extractable quantity of sedimentary DNA (μg g^−1^ sediment) decreased from approximately 10 μg g^−1^ at 1 cm to ~7.5 μg g^−1^ at 10 cm (Figure 2E). Assuming that DNA pools are approximate to microbial biomass, these values infer that microbial abundances decrease with depth, which is consistent with observations in both shallow and deep-sea sediments (Kallmeyer et al., 2012; Chen et al., 2017). Higher microbial biomass near the sediment-water interface is consistent with this being the zone where the most labile organic matter is deposited at the sediment surface, while organic matter becomes more refractory with depth. However, macrofauna may confound these interactions by actively exchanging and flushing bottom water into deeper sediments, thereby introducing more labile organic matter deep into the sediments. Across the season, DNA pools were distinct between fall and winter months versus summer months (Figure 2F). These variations may be caused by fall phytoplankton blooms that are stored in the sediments over the winter months, and decrease in the summer months due to the activity of microorganismal mineralization of detrital organic matter, including degradation and metabolism of exogenous DNA. This is evident when sedimentary DNA pools are plotted as a function of temperature, which is negatively correlated; as temperature increased, DNA pools decreased (Figure 3C).

### Iron cycling microorganisms

There were OTUs that could be unambiguously assigned to be involved in the iron biogeochemical cycle. The iron-oxidizing Zetaproteobacteria formed an OTU (Otu00040) that was 0.28% median seasonal abundance (range 0.01-2%), which is a typical relative abundance for this class of bacteria in marine sediments (Rubin-Blum et al., 2014; McAllister et al., 2015; Laufer et al., 2016, 2017; Beam et al., 2018). The Zetaproteobacteria relative abundances changed with respect to depth (Figure 4A), but did not exhibit pronounced seasonal variation except in one sample in September that appeared to correlate with increased Fe_HR_ production (Figure 4B; Figure 2D). The Zetaproteobacteria appear to be confined to the upper 5 cm of sediment, and have the highest relative abundances around 3-4 cm (Figure 4A), which has been previously documented from a depth profile of 16S rRNA genes from sediment and iron oxide macrofaunal burrow walls at this mudflat (Beam et al., 2018). The peak abundance of Zetaproteobacteria also matches well with the peak in porewater Fe(II) and Fe_HR_ from these sediments (Figure 2A, C), which suggests that they are most abundant in the active iron cycling zone—perhaps a common pattern for Zetaproteobacteria in bioturbated sediments (Beam et al., 2018). Although the Zetaproteobacteria exhibit a relatively low median abundance, our previous work has shown their abundance increases significantly in iron-oxide coated worm burrow walls, where they are likely involved in Fe(II) oxidation coupled to oxygen reduction and carbon dioxide fixation (Beam, 2018; McAllister et al., 2015). Consistent with these observations is that increased dark CO_2_fixation with concomitant Fe(II) oxidation has been observed in burrow walls of the polychaete *Neries diversicolor* (Reichardt, 1986), which was the most abundant polychaete at the Eddy. Thus, dark CO2 fixation by iron-oxidizing Zetaproteobacteria may contribute to the autochthonous organic matter pools in coastal sediments, similar to that observed in autotrophic sulfur-oxidizing Gammaproteobacteria (Dyksma et al., 2016).

**Figure 4.**
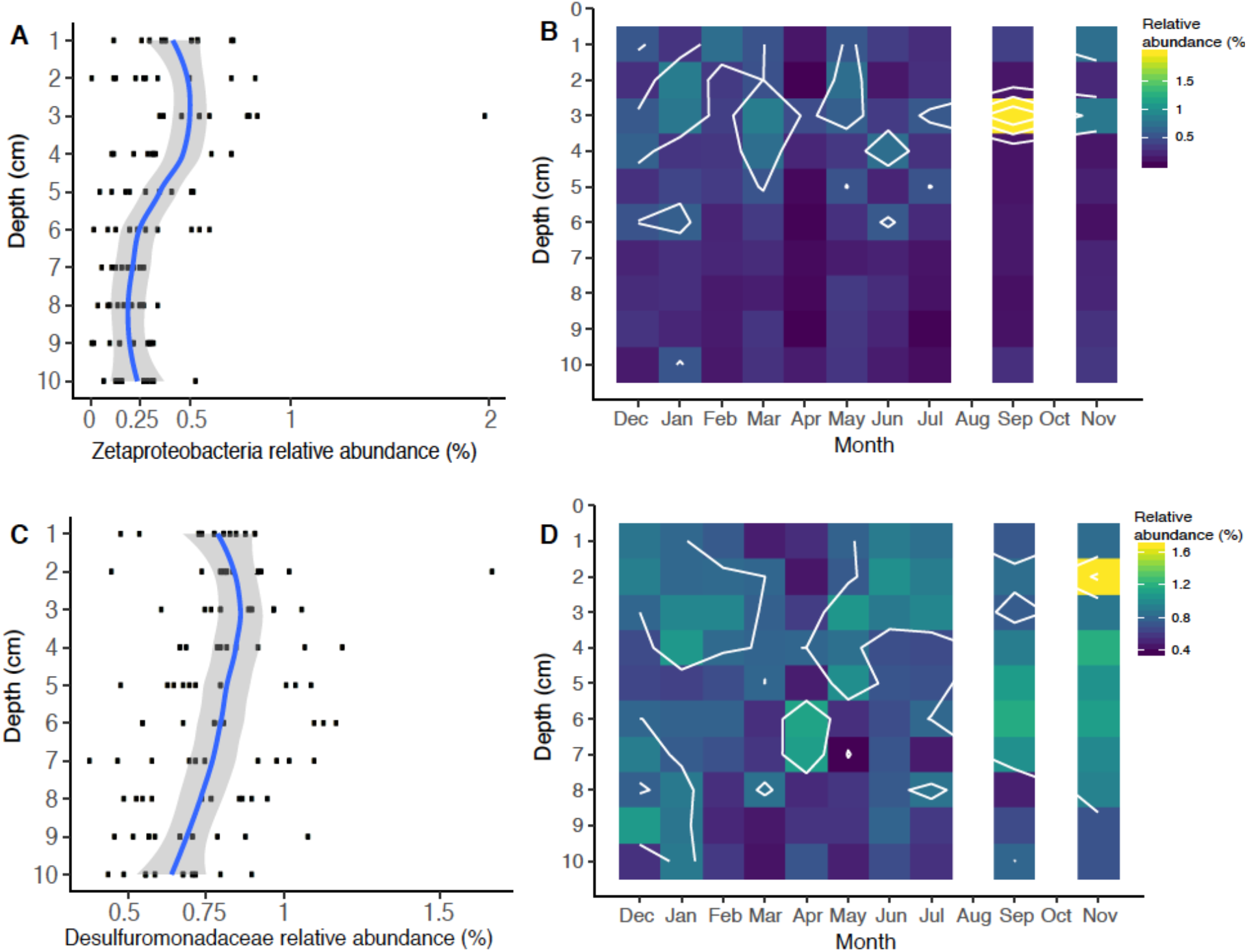
Relative abundance of iron cycling operational taxonomic units (OTUs) identified over the season: Iron-oxidizing Zetaproteobacteria (OTU00040) relative abundance as a function of depth (**A**) and season (**B**), and putative iron-reducing Desulfuromondaceae (OTU00028) as a function of depth (**C**) and season (**D**).

Microorganisms that catalyze the reduction of iron are phylogenetically diverse (Melton et al., 2014) and, particularly in marine sediments, are more difficult to identify with 16S rRNA gene profiling. However, culture-based studies from marine sediments have suggested that members of the Deltaproteobacteria family Desulfuromonadaceae can reduce iron, as well as other some other Deltaproteobacteria lineages (Vandieken et al., 2006), which have been subsequently identified in coastal sediments (Reyes et al., 2017; Otte et al., 2018; Buongiorno et al., 2019). We identified one OTU containing members of the Desulfuromonadaceae family (Otu00028) that was relatively abundant across the season accounting for a median relative OTU abundance of 0.80% (range=0.40-1.7%) (Figure 4C, D). The Desulfuromonadaceae OTU was also most abundant in the active iron cycling zone (Figure 6C), which appears to mirror the porewater and highly reactive iron profiles (Figure 2A, C). The Desulfuromonadaceae abundances fluctuated over the season (Figure 4D) perhaps in response to organic matter and/or iron oxide availability. Correlations of Desulfuromonadaceae with porewater and highly reactive iron geochemistry were not indicative of a strong relationship (not shown), but this does not preclude a causal relationship. Canonical iron-reducing bacterial genera such as *Geobacter* and *Shewanella-like* OTUs were extremely rare or below detection in these sediments, similar to findings from coastal sediments near Denmark and Sweden (Reyes et al., 2017; Otte et al., 2018; Buongiorno et al., 2019).

The relative abundances of Zetaproteobacteria and Desulfuromonadaceae were low compared to other OTUs. A primary explanation for this is that the most active iron cycling may occur in association with burrow walls, which only make up a small percentage of the total area we sampled, due to sampling at a low spatial resolution. Accordingly, it is possible their abundances are relatively high in the hotspots of active iron cycling, indicating that they are very active in iron cycling in marine sediments despite their low overall abundances. This also suggests that although their abundance is low, these iron cycling microorganisms can profoundly affect the porewater Fe(II) and Fe_HR_ pools across a large spatial gradient. There may also be undescribed microbial lineages involved in both the oxidation and reduction of iron. These would be unaccounted for based on our OTU-based taxonomic analysis. Nevertheless, iron cycling microorganisms are observed throughout the season, and catalyze essential biogeochemical transformations in these environments.

### Seasonal dynamics of the sedimentary microbial community

The sedimentary microbial communities were dominated by bacteria (total OTU abundance = 97.5%), with minor contributions by archaea (total OTU abundance = 2.5%)—consistent with the observations of Chen et al. (2017) and Kim et al. (2008) that bacteria dominate over archaea in bioturbated sediments. The three most abundant bacterial OTUs throughout the season were related to the Deltaproteobacteria and one was associated with the Gammaproteobacteria (Figure 5). Deltaproteobacteria and Gammaproteobacteria are commonly observed as abundant members of coastal sediment microbial communities (Kim et al., 2008; Lasher et al., 2009; Wang et al., 2012; Chen et al., 2017; Cleary et al., 2017; Qiao et al., 2018). The two Deltaproteobacteria OTUs were related to the families Desulfobulbaceae (Otu00001; median seasonal abundance=6.4%; range=3.0-8.8%) and the genus Desulfococcus in the family Desulfobacteraceae (Otu00002; median seasonal abundance=5.7%; range=1.6-9.8%), which are likely involved in sulfur oxidation and sulfate reduction, respectively, in these sediments. The Desulfobulbaceae is a diverse family within the Deltaproteobacteria that contains the so-called ‘cable bacteria’ that transport electrons from sulfur oxidation over centimeter scale distances and catalyze oxygen reduction at sediment-water interfaces (Nielsen and Risgaard-Petersen, 2015). We have observed on many occasions the presence of cable bacteria-like filaments from this field site (Figure 1F), and hypothesize that they are functional in these bioturbated sediments, which has been recently demonstrated (Aller et al., 2019). Cable bacteria have been observed to contribute to both iron and sulfur cycling in static marine sediments (Nielsen et al., 2010; Meysman et al., 2015; Nielsen and Risgaard-Petersen, 2015; Otte et al., 2018) and are also likely impacting the iron and sulfur cycles in bioturbated sediments (Burdorf et al., 2017; Aller et al., 2019).

**Figure 5.**
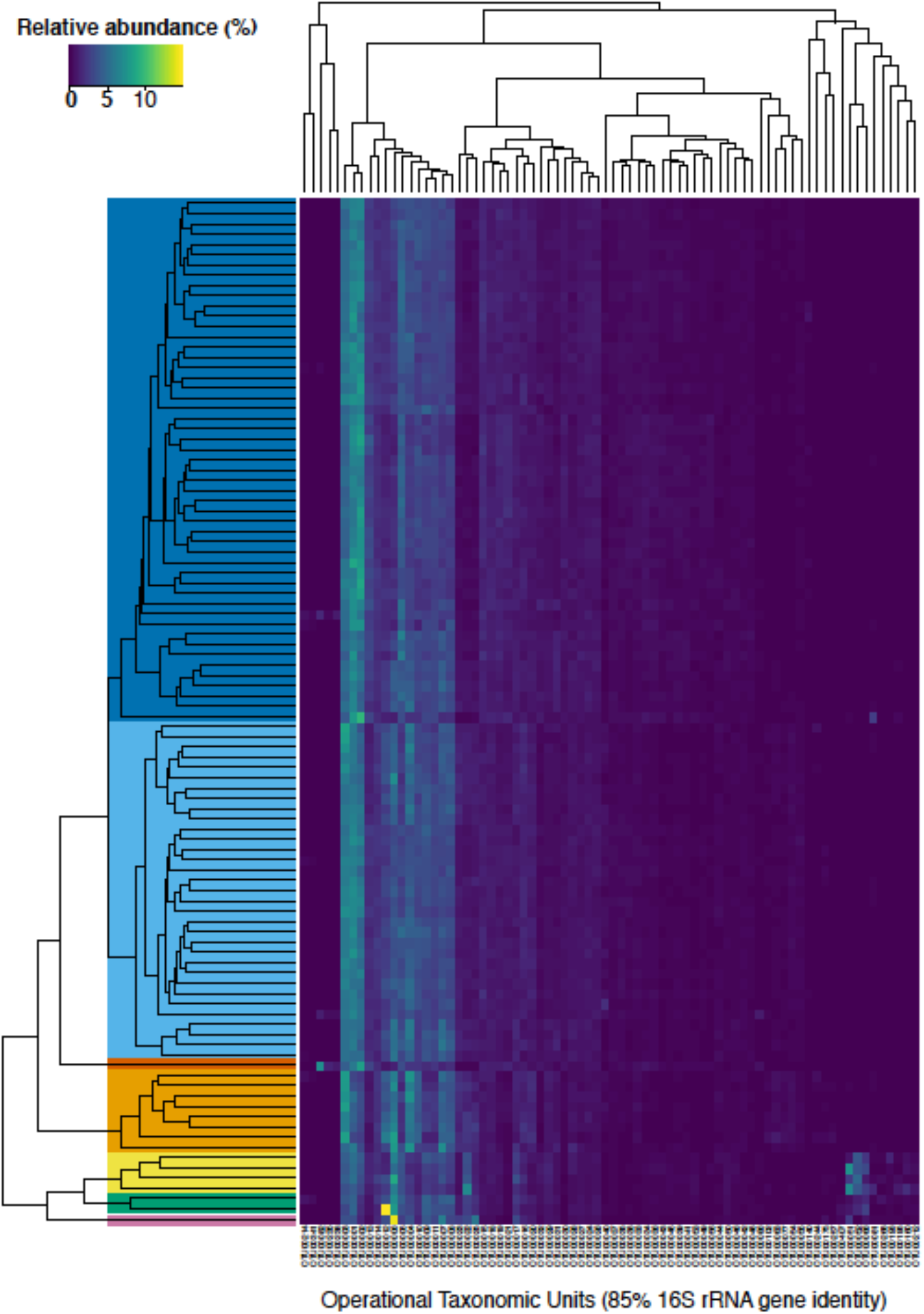
A heatmap of the relative abundance of bacterial and archaeal taxonomic units (OTUs) clustered at 85 % identity that were at least 0.5 % relative abundance in one sample (n=76 OTUs). The different samples were clustered using average linkage hierarchical clustering of the Bray-Curtis dissimilarity as implemented in the R package vegan (Oksanen, 2019). Solid colors depict the different ecostates that were selected from the clustering analysis (see Figure 5). OTU identifiers can be referenced with specific taxonomic groups in Table S1.

Each discrete spatiotemporal sample (n=100) was clustered based on the relative abundance of operational taxonomic units (OTUs) at 85% 16S rRNA gene identity. The composition of the sedimentary microbial community did not exhibit large fluctuations with the changing seasons in this bioturbated mudflat (Figure 5, 6). Although some individual OTUs exhibit spatial and temporal variability, the community was generally comprised of approximately 20-25 dominant OTUs (Figure 5). Seven main clusters predominated the various communities with respect to time and space (Figure 5). These clusters comprise different ‘ecostates’ that were identified at various depths and times during the season. Ecostates are operationally defined as a collection of inferred taxonomic groups from distinct ecological samples, and they can be used to track changes in ecological communities through space and time (O’Brien et al., 2016). There is a strong depth stratification of ecostates that persists throughout the season (Figure 6). Ecostate 2 typically dominates all depth intervals < 5 cm over the season, which is likely a result of the higher oxygen concentrations in shallower sediments due to surface diffusion and enhanced biodiffusive transport by animals (Glud, 2008). Ecostate 1 was usually confined to deeper sediments > 5 cm where lower oxygen conditions would be prevalent (Glud, 2008) (Figure 6). The presumed presence of labile organic matter in the upper 5 cm of sediment also likely contributes to the separation of ecostates 1 and 2 as has been observed between other surface and subsurface coastal sediments (Cleary et al., 2017; Qiao et al., 2018). These ecostates are relatively stable throughout the season (Figure 6), and appear to be both resistant and resilient to the observed seasonal changes in environmental conditions and other stochastic events likely occurring in the intertidal zone. More than half of the samples at the sediment-water interface were classified as ecostate 3, which suggests some unique properties of this interface that contributes to the observed community signal (Figure 5, 6). This consistent pattern of stratification was only disrupted once, in the April profile. This profile differed markedly, with three unique ecostates (4, 5, and 6) that were not observed in any other month. These ecostates contained a cluster of high relative abundances of OTUs, described below, that only comprise a minor community component of the other ecostates (Figure 4). The occurrence of unique ecostates in April could reflect a community response to ecosystem disturbance or a stochastic assembly of horizontal spatial heterogeneity, which would both imply resilience of the system.

**Figure 6.**
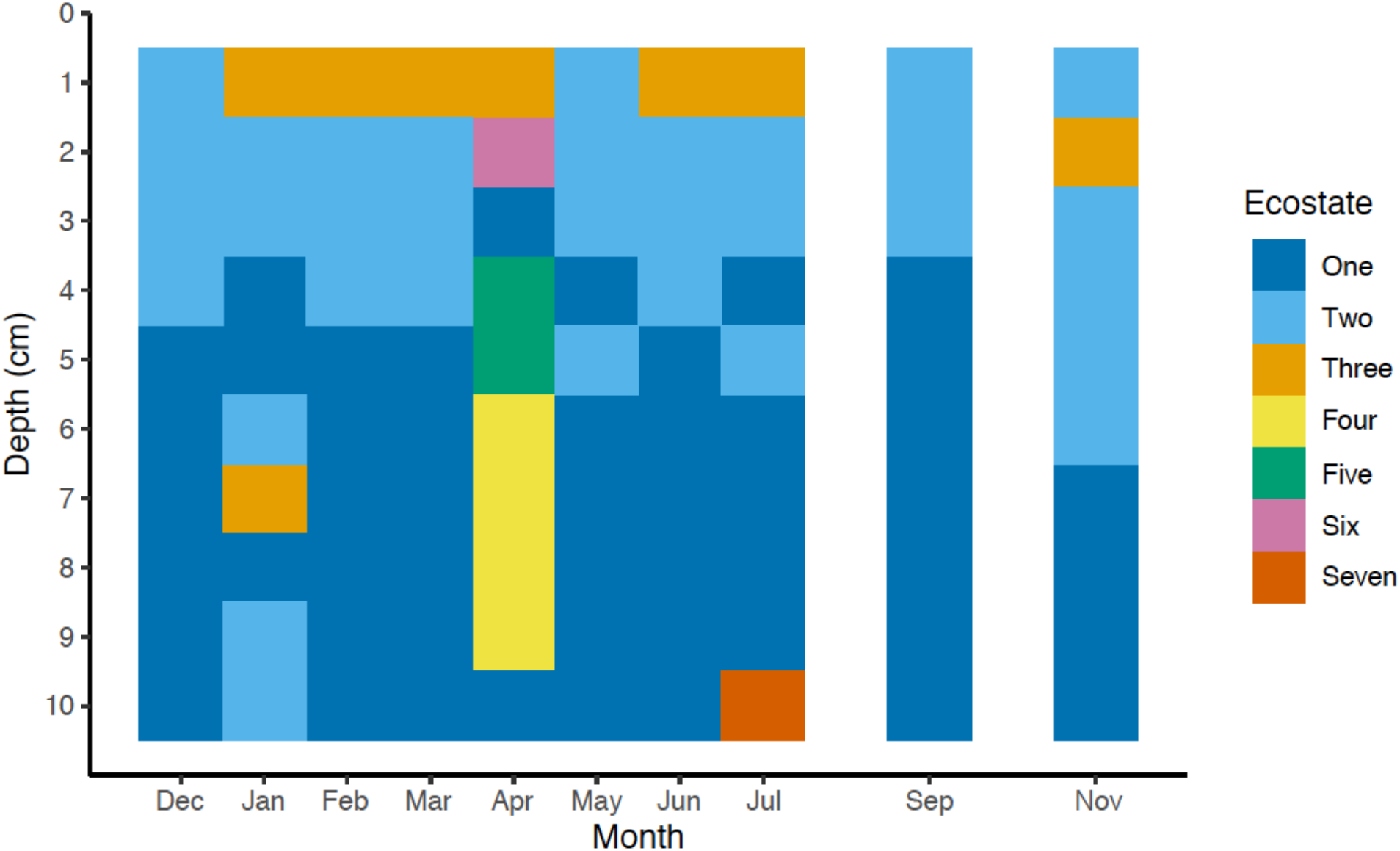
Ecostates (n=7) of bacterial and archaeal microbial communities across one seasonal cycle. The seven different ecostates were identified from the Bray-Curtis dissimilarity matrix of the relative abundance of all OTUs (n=12,437) by average linkage hierarchical clustering. Taxonomic assignments of all taxa with a relative abundance ≥ 1.0% for each ecostate are shown in Table S2.

Approximately 50% of the taxa assigned to each of the different ecostates, either at the order, family, or genus level, could be associated to known physiological representatives that are presumed to be either predominately anaerobic or aerobic. For example, all known relatives of the family Desulfobulbaceae, or genus Desulfococcus, predominant in Ecostates 1 and 2 are known to be anaerobes; while known relatives of the family Flavobacteriaceae, and family Marcinicellaceae (both abundant in ecostate 3) are obligate aerobes. From this we surmise that the overall dominant ecostates 1 and 2 had relatively high abundances of putative anaerobes, compared to putative aerobes (Supplementary Table S2), while ecostate 3, found primarily in the top 1 cm (Fig 5) had higher relative abundances of putative aerobes, and corroborates the availability of oxygen declining with depth (Glud, 2008). In general, we found members of the phyla Bacteroidetes more prevalent in ecostates 2, 3, and 6, all associated the upper 5 cm of the sediments than in deeper ecostates 1 and 7. The Bacteroidetes are known for their involvement in the degradation of more complex organic matter, either aerobically or anaerobically (Kirchman, 2002; Hahnke et al., 2016). This is consistent with the most labile organic matter being present in the upper layers of sediment, in part driven by spring and fall phytoplankton blooms. Ecostates 4, 5, and 6 were only present in April, and OTUs that increased in abundance were Flavobacteriaceae (Otu00006), Rhodobacteraceae (Otu00022), Vibrionales (Otu00051), Helicobacteraceae (Otu00052), unclassified Bacteriodetes (Otu00009), Clostridiales (Otu00066), and Alteromonadales (Otu00012) OTUs (Table S1). These OTUs may be involved in the degradation of phytoplankton-derived organic matter—relatives of some of these OTUs have been observed to change abundance in response to phytoplankton-derived organic matter as well as plant-derived organic matter (Raulf et al., 2015). OTUs that were elevated in April were similar to the elevated OTUs in ecostate 3, which further supports the hypothesis that these OTUs are potentially involved in organic matter-specific responses such as carbon mineralization. Although these microbial responses to phytoplankton blooms are tentative, it would be worth exploring in other coastal environments with respect to seasonal variation.

### Seasonal dynamics of eukaryotic phytoplankton in sediments

An important ancillary question related to our focus on iron-cycling communities, is the role for pelagic-derived phytoplankton, such as diatoms, to be transported to these intertidal sediments by sedimentation and/or potentially by the activity of suspension feeding macrofauna. These phytoplankton may act as an important source of labile organic matter in sediments at different times of the season (Kristensen, 2001; Kristensen and Mikkelsen, 2003). To address this, we used 18S rRNA gene sequencing to determine the presence and relative abundance of phytoplankton and other eukaryotes in these sediments over the seasonal cycle. We recovered 18S rRNA gene amplicons from 99/100 samples over the season from depths of 1-10 cm. However, the total number of quality filtered sequences from each sample varied significantly with a median of 2,636 sequences per sample and a range of 1-24,484 sequences. One explanation for this variability is that it reflects biological reality, in that there are simply few eukaryotes present in those low-yield samples, thus yielding few 18S rRNA gene sequences. Pasulka et al. (2019) utilized a similar approach to investigate benthic microbial eukaryotes at Guaymas Basin hydrothermal vents, and also recovered a highly variable numbers of sequences among samples. Further sequencing of samples from marine benthic habitats will be needed to understand the relationship between sequence yields, the biomass of microbial eukaryotes in unprocessed samples, DNA template concentrations in extracted samples, and possible PCR inhibition in these complex and heterogeneous environmental DNA samples. It cannot be ruled out that some sequences are sourced from relict, extracellular DNA bound to sediment particles, and not indicative of the conditions that prevailed during the course of this study. However, a recent paper looking at marine sediments found little influence of extracellular DNA on amplicon studies similar to these samples (Ramírez et al., 2018). To further reduce the confounding role of extracellular DNA, we focused only on relatively abundant diatom taxa (>0.1% of the total eukaryotic community), as discussed below.

With these caveats in mind, we found that diatoms (Bacillariophyta) comprised a median seasonal relative abundance of 9.5% with a range of 0-33.5% of all 18S rRNA gene sequences, and followed distinct spatiotemporal patterns (Figure 7). Their abundance appeared to follow a seasonal pattern, with small increases in abundance occurring in the winter months of January and February, and also in the fall months of September and early November (Figure 7). These increases in diatom relative abundance may coincide with spring and fall phytoplankton blooms occurring in the water column. Although, in shallow estuaries along the Maine coast, primary productivity by phytoplankton may occur year-round, and not follow discrete seasonal succession as observed in other, deeper estuary and coastal waters (Thomas et al., 2003). We hypothesized that the activity of burrowing and irrigating animals would transport phytoplankton deep into these intertidal sediments. Animals in this mudflat, in particular *N. diversicolor*, are known to feed on phytoplankton throughout the season, and may increase feeding in response to increased food availability during phytoplankton blooms (Riisgard, 1991; Christensen et al., 2000; Kristensen, 2001). In some of the months (that is, December, January, and November), diatoms were observed at the deepest sampling depth of 10 centimeters (Figure 7), which we view as an indication of active transport by irrigating animals in combination with physical settling of diatom blooms in mid-winter and fall seasons. Animals may burrow deeper in winter months as well providing a conduit for pelagic diatoms to be transported to deeper sediments; in the summer, animals may produce shallower burrows, and thus phytoplankton would remain closer to the sediment-water interface (Esselink and Zwarts, 1989).

**Figure 7.**
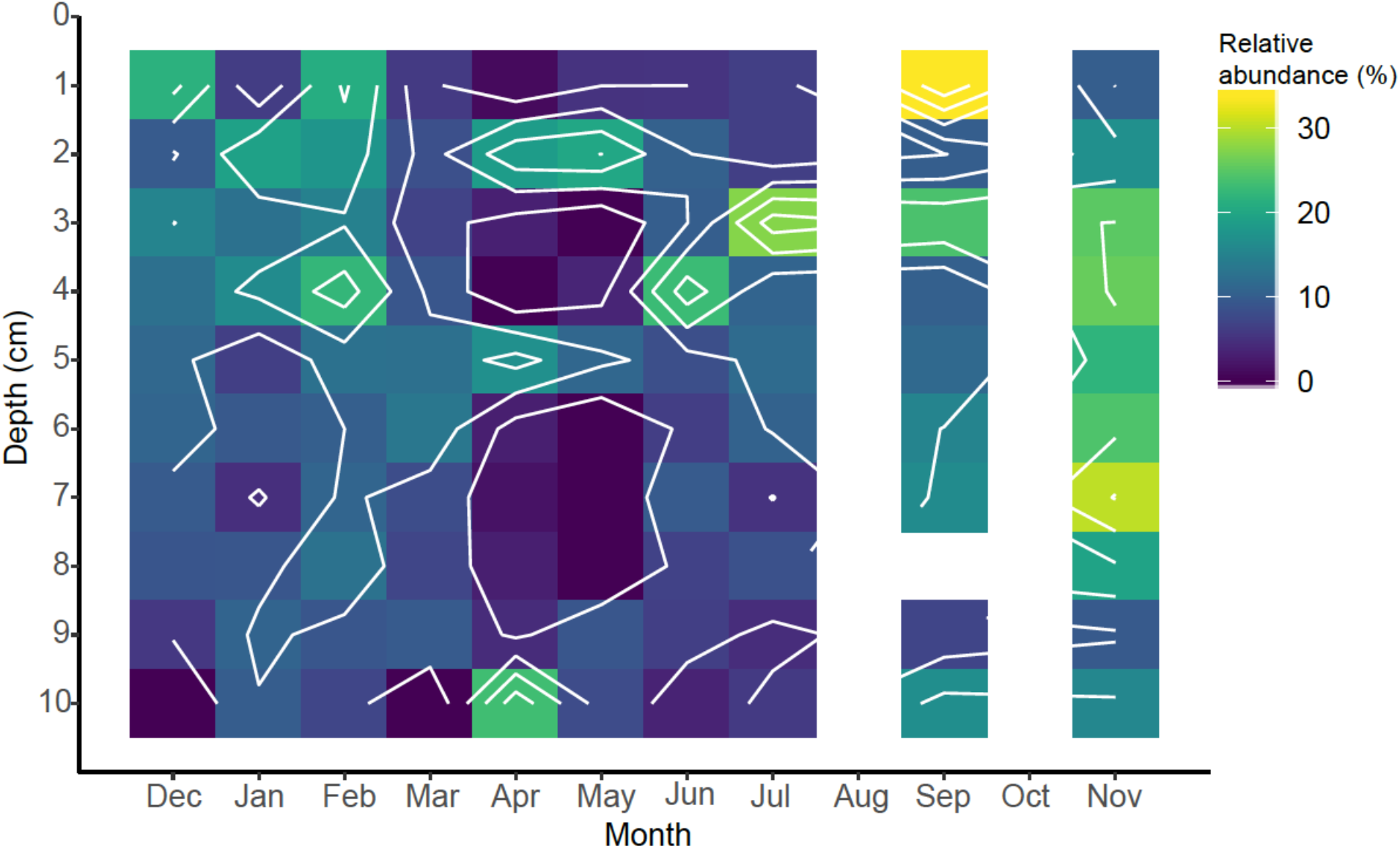
Spatiotemporal relative abundance of diatoms (Bacillariophyta) in sediments identified by 18S rRNA gene sequencing.

A total of 119 eukaryotic OTUs were present at a relative abundance of ≥ 0.1% of total sequences (range=0.10-3.87%; Table S4). Of these OTUs, 34 were identified as belonging to the two most abundant classes of eukaryotic marine phytoplankton, including the diatoms (n=18 OTUs) and dinoflagellates (n=16 OTUs). Presence of either of these phytoplankton groups in subsurface sediments could indicate their transport via burrowing-based ventilation, or preservation via sedimentation and burial over substantially longer periods of time. Many of the most abundant phytoplankton OTUs were associated with benthic habitats (Table S4). Diatoms and dinoflagellates were identified as four of the ten most abundant OTUs and included a dinoflagellate from the order Peridiniales (OTU0004, 3.87%), a raphid-pennate diatom (OTU0006, 3.24%), the diatom *Psammodictyon* sp. (OTU0009, 2.85%) and the dinoflagellate *Islandinium minutum* (OTU0010, 2.72%). The Peridiniales include a diverse group of dinoflagellates rendering the taxonomic designation of OTU0004 somewhat ambiguous. Nonetheless, OTU0004 is notable with respect to its prevalence across depth, with greater proportions found deeper in the sediments. Raphid-pennate diatoms include both pelagic and benthic species, but the motile benthic species are thought to be substantially more diverse than pelagic types (Mann et al., 2017), and are likely the type associated with OTU0006. The diatom *Psammodictyon* (OTU0009) is also associated with benthic habitats, living at the sediment water interface (Round et al., 1990). *Islandinium minutum* (OTU0010) is primarily associated with a particular ‘spiny’ morphology of dinoflagellate cysts that are common in high latitude sediments (Potvin et al., 2013). Several other cyst forming dinoflagellates were detected in the next five most abundant OTUs (including OTU0011, 12, 14), and were identified as the ‘red tide’ bloom forming species *Alexandrium fundyense* (OTU0011 and 14) and *Woloszynskia halophila* (OTU0012). *Alexandrium fundyense* is well known to coastal Maine, where it causes harmful algal blooms (HABs) that subsequently encyst and overwinter in the sediments (Anderson et al., 2014) while *Woloszynskia halophila* forms blooms and abundant cysts in the Baltic Sea (Kremp et al., 2008). The extent to which phytoplankton distributions in the sediments can be used as indicators of subsurface ventilation remains to be tested under more controlled experimental conditions, but the possibility exists that these organisms could serve as useful biomarkers for this important biogeochemical process.

### Sedimentary macro-, meio-, and micro-fauna

The 18S rRNA sequences also revealed the presence of *Nereis (Hediste) diversicolor*, which was the presumed dominant polychaete worm in this mudflat. This was the most abundant polychaete 18S rRNA gene sequence, accounting for 3.7% of the total 18S rRNA gene sequences (n=14,149 sequences). Mollusca 18S rRNA gene sequences were not a very abundant component of any samples, although we detected 0.04% of the total 18S rRNA gene sequences (n=148 sequences) related to the small, intertidal-dwelling clam, *Macoma balthica*, which is common to the Sheepscot Estuary and to North Atlantic intertidal sediments (Stickney, 1959). We have observed small clams similar to *M. balthica* from the Eddy on numerous occasions.

Nematodes can comprise a large fraction of the biomass of marine sediments (Heip et al., 1985) and are bewilderingly diverse (Bik et al., 2010). The 18S rRNA gene libraries revealed that nematodes comprised a median relative abundance of 21.9% with a range of 0-67.2% of the total 18S rRNA gene sequences. Their relative abundance was highest near the sediment-water interface, and abundance declined with depth (Figure 8). This finding corroborates observations primarily based from direct species counts and identification (Montagna et al., 1989). Nematode abundance and grazing activity have also been observed to increase bacterial activity and diversity in terrestrial sediments (Jiang et al., 2017). Nematodes play an important role in grazing of bacteria and other microbial eukaryotes in sediments (Pascal et al., 2008), and potentially play an important role in controlling iron biogeochemical dynamics.

**Figure 8.**
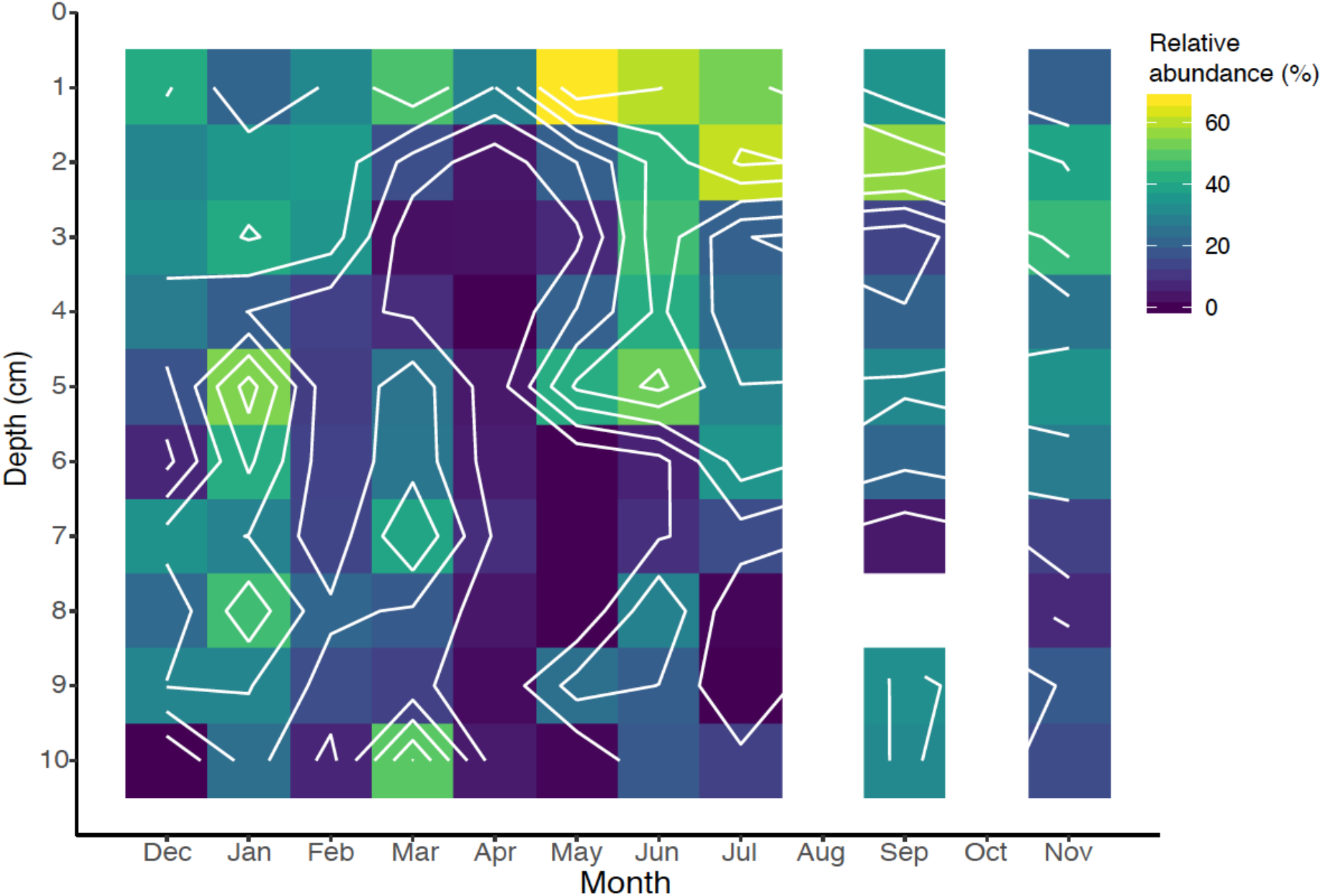
Spatiotemporal relative abundance of nematodes (Nematoda) identified by 18S rRNA gene sequencing over the seasonal cycle.

Protozoa can also comprise a significant portion of the diversity in sedimentary microbial communities (Fenchel, 1969; Forster et al., 2016), and their median seasonal abundance of all 18S rRNA gene sequences was 57% (range=17.3-100%). Similar to the nematodes, the protozoan communities also exhibited pronounced seasonal peak abundances in summer months most likely in response to increased microbial food availability (Figure 9). The sedimentary protozoa community was enriched in the deeper part of the sediment (Figure 9), which might imply that they are active in this mudflat in anoxic carbon mineralization (Fenchel, 1978). Protozoan grazing may also influence the abundance and activity of bacteria and archaea in marine sediments (Starink et al., 1994), and would play a currently unknown role on the influence on the sedimentary iron biogeochemical cycle.

**Figure 9.**
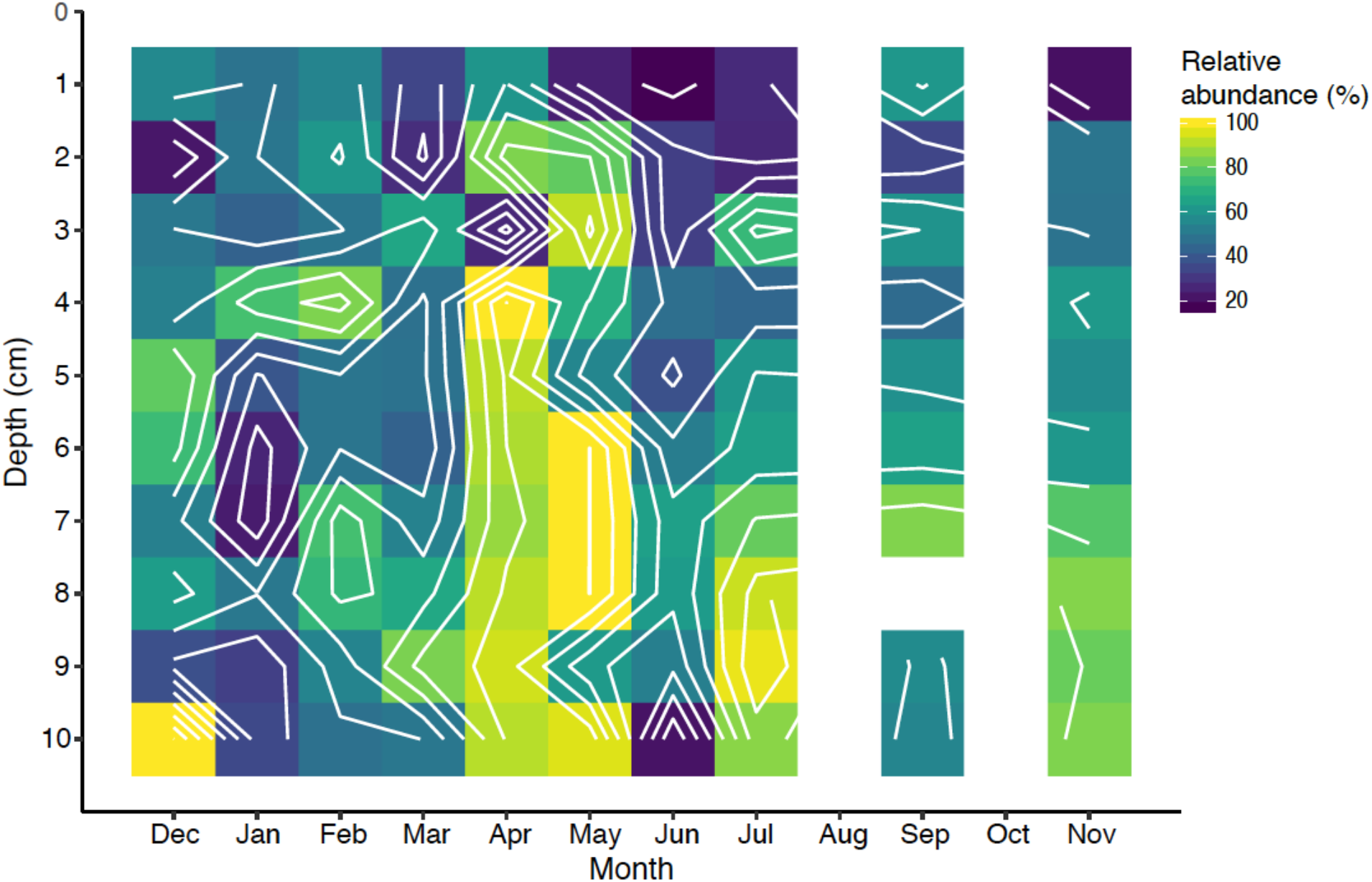
Spatiotemporal relative abundance of protozoa identified by 18S rRNA gene sequencing over the seasonal cycle.

## Concluding remarks

We observed spatial and temporal heterogeneity in both porewater dissolved Fe(II) and highly reactive Fe (Fe_HR_) sedimentary pools over the seasonal cycle that are primarily controlled by the master variable, temperature. We infer that changes in temperature influence both animal feeding and irrigational behavior (Kristensen, 2001; Fang et al., 2019) as well as the direct influence of temperature on increased microbial iron oxidation and reduction. Thus, the timing and spatial extent of macrobiota-microbiota relationships can provide information about the potential increase or decrease of iron biogeochemical transformations in coastal sediments. Understanding these macrobiota-microbiota relationships in greater detail could shed light on important ecosystem processes such as seasonal changes in Fe fluxes from coastal sediments, which are a major source of bioavailable Fe to the ocean. The sedimentary microbial communities were fairly stable throughout the seasonal cycle with a major demarcation between surface and subsurface sediments. Any perturbation in these microbial community ecostates was rapidly recovered in the following sampling period, reverting to the steady-state ecostate conditions, which could be an internal functional resilience mechanism of these communities to natural environmental changes. The sediment mixing and irrigational behavior of bioturbating animals may be a contributing factor to this microbial community resilience. The presence and quantification of phytoplankton in these bioturbated sediments throughout the season may provide clues about changes in macrofaunal feeding behavior and potential impact on the iron biogeochemical cycle. Our observations also suggest that specific benthic micro- and meiofauna associations or interactions likely play an, as yet poorly understood, role in controlling the sedimentary iron biogeochemical cycle. In the rapidly changing coastal ocean system, it is imperative to conduct both temporal and spatial monitoring of sedimentary ecosystems, as environmental perturbations could be reflected in the microbial communities and the essential biogeochemical transformations they catalyze.

## Acknowledgements

Primary funding for this research was provided by a National Science Foundation grant OCE-1459600 to DE, PRG, and DTJ. Additional support came from NSF OIA-1849227 to DE, NRR, and PC., and from NASA grant NNX16AG59G, and Bigelow Laboratory institutional funds. JPB thanks Dr. Peter Larsen for many helpful and insightful discussions about benthic macrofaunal behavior and identification, and Dr. Jarrod Scott for inspiring the microbial community sampling design and insightful discussions on microbial community analysis across spatial and time series data. We also thank Dr. Alex Michaud for insightful comments on iron and sulfur sedimentary biogeochemistry, and Dr. Jeremy Rich for helpful comments on the manuscript. We appreciate the assistance of organizing and shipping DNA samples for sequencing by Jennifer Delany at Harvard University.

## Supplemental Table Captions

**Table S1.** Seasonal physicochemical data.

**Table S2.** The top 76 OTUs for each ecostate in order from the highest relative abundance to the least relative abundance.

**Table S3.** This table is derived from the different ecostate taxa counts from Table S1. For each ecostate, all taxa of ≥1% relative abundance are represented. Taxa are classified to the highest taxonomic resolution against the Silva database, along with either their phylum, or in the case of the Proteobacteria, their corresponding class, abbreviated as Delta, Gamma, Alpha, or Epsilon. The colors denote whether taxa were judged to be presumptive anaerobes (red), possible anaerobes (purple), or presumptive aerobes (green). These physiological categories were assigned based on multiple cultured representatives of family, order, or class being known anaerobes or aerobes. For approximately half the taxa there were not enough cultured relatives known to make any judgement. For each ecostate, the % of Gammaproteobacteria, Deltaproteobacteria, and Bacteroidetes are given, since these were the three most abundant taxa within all the ecostates. The % anaerobes and % aerobes are the sums of the relative abundances of presumptive and possible anaerobes, or presumptive aerobes for each ecostate.

**Table S4.** Eukaryotic operational taxonomic units (OTUs) and taxonomic classifications from all seasonal samples.

